# Effects of calorie restriction on reactive oxygen species production by mitochondrial reverse electron transport, mitochondrial permeability transition pore, and beta-adrenergic stimulation during cardiac hypertrophy

**DOI:** 10.1101/2022.02.02.478846

**Authors:** Aline Maria Brito Lucas, Plinio Bezerra Palacio, Pedro Lourenzo Oliveira Cunha, Heberty Tarso Facundo

**Affiliations:** Faculdade de Medicina, Universidade Federal do Cariri, Barbalha, CE, Brazil

**Author notes:** Corresponding author: Dr. Heberty Tarso Facundo, Universidade Federal do Cariri, Rua Divino Salvador, 284, Barbalha, Ceará, Brazil, 63180-000.

**Keywords:** Mitochondria, calorie restriction, cardiac hypertrophy, oxidative stress, adrenergic signaling

## Abstract

Calorie restriction is a nutritional intervention that reproducibly protects against the maladaptive consequences of cardiovascular diseases. Pathological cardiac hypertrophy leads to cellular growth, dysfunction (with mitochondrial dysregulation), and oxidative stress. The mechanisms behind the cardiovascular protective effects of calorie restriction are still under investigation. In this study, we addressed the impact of calorie restriction on mitochondria, oxidative stress markers, and β-adrenergic signaling during cardiac hypertrophy. This dietetic intervention prevented cardiac protein elevation, decreased atrial natriuretic peptide levels, and blocked the increase in heart weight per tibia length index seen in isoproterenol-induced cardiac hypertrophy. Our data suggest that inhibition of cardiac pathological growth by calorie restriction is accompanied by a lower mitochondrial reactive oxygen species formation and improved mitochondrial content. We also found that superoxide dismutase and glutathione peroxidase activities negatively correlate with cardiac hypertrophy. Calorie restriction also attenuated the opening of the Ca^2+^-induced mitochondrial permeability transition pore in mitochondria isolated from isoproterenol-treated mice. Isoproterenol (a β-agonist) increases cardiac rate (chronotropic response) and force of contraction (inotropic response). Given the nature of cardiac hypertrophy induction by isoproterenol, we tested whether calorie restriction could change the cardiac β-adrenergic sensitivity. Using isolated rat hearts in a langendorff system, we found that calorie restriction mice (similar to controls) have preserved β-adrenergic signaling. On the flipside, hypertrophic hearts (treated for seven days with isoproterenol) were insensitive to β-adrenergic activation using isoproterenol (50 nM). Despite protecting against cardiac hypertrophy, calorie restriction did not alter the lack of responsiveness to isoproterenol of isolated hearts harvested from isoproterenol-treated rats. These results suggest (through a series of mitochondrial, oxidative stress, and cardiac hemodynamic studies) that calorie restriction possesses beneficial effects against hypertrophic cardiomyopathy. However, it may lack effects on some of the hypertrophic consequences, such as β-adrenergic signaling repression.

## 1. INTRODUCTION

Cardiac hypertrophy is the enlargement of fully differentiated cardiomyocytes which is accompanied by sarcomere reorganization, enhanced protein synthesis, and associated reactivation of the fetal gene program. Cardiac hypertrophy is considered a compensatory adaptation but, if sustained, will lead the myocardium to functional and histological deterioration. This attempt to compensate for dysregulations will normalize higher wall tension of the ventricles but will lead to heart failure. Known inducers of cardiac hypertrophy such as angiotensin II, alpha and beta-adrenergic agonists, tumor necrosis factor-alpha, ouabain, or cyclic stretch have all been characterized as intracellular Reactive Oxygen Species (ROS) inducers. (1–6). ROS are considered pro-hypertrophic molecules that mediate hypertrophic signals leading to cardiac hypertrophy/heart failure. Instead, inducing antioxidants protect or avoid this pathological growth (7–9). The involvement of ROS and the beneficial effects of antioxidants in pathological cardiomyopathy highlights the need for a better understanding of the mechanisms. Strategies aimed at blocking cardiac hypertrophy by lowering oxidative stress would be of great value. One attractive dietetic tool candidate for cardiac hypertrophy/heart failure treatment is calorie restriction which, can be easily induced and has been shown to protect against cardiac pathologies, including cardiac hypertrophy (1, 10–12).

The limitation of calorie ingestion in the absence of malnutrition (the so-called calorie restriction) protects against cardiovascular diseases besides extending lifespan (12). Calorie restriction protects cardiac tissue from hypertrophy development (10). Calorie restriction also avoids cardiac inflammation, fibrosis and protects against cardiac iron dysregulations in ob/ob and db/db mice [13]. This dietetic intervention ameliorates the diastolic dysfunction and remodeling induced by metabolic syndrome in obese rats (11). Avoiding excessive mitochondrial or tissue ROS production is part of the mechanism which will impact the oxidative modifications of lipids, DNA, and proteins. Calorie restriction is a powerful non-pharmacological tool that can avoid the array of detrimental consequences resulting from oxidative stress (1, 14–17). Additionally, it counteracts sympathetic signaling dysregulations (18). Indeed, the sympathetic system activity increases during aging (19). An autonomic imbalance (higher sympathetic and reduced parasympathetic) may play a major role in cardiomyopathy/heart failure (20). This event is especially important during aging.

Although the mechanisms by which calorie restriction protects against cardiovascular diseases are multifactorial and poorly understood, mitochondria seem to be involved in the cellular adaptations leading to its effects. As master regulators of cellular metabolism generating ATP and ROS, these organelles are attractive for strategies aiming at preserving cardiomyocyte function. Mitochondria occupies about 30% of the cardiomyocytes’ mass. Furthermore, their activity correlates with oxygen consumption and heart rate. These organelles consume largely fatty acids and provide more than 95% of the ATP used by the heart. Mitochondria are also involved in several intracellular signaling, cell death, and calcium homeostasis (21). Calcium overaccumulation into mitochondria leads to the opening of the mitochondrial permeability transition pore (mPTP). mPTP triggers a sudden increase in inner mitochondrial membrane permeability with consequent swelling (22). This phenomenon is particularly important since swelling secondary to mPTP leads to rupture of mitochondrial membranes promoting cell death by necrosis or apoptosis. In cardiomyocytes, mitochondrial Ca^2+^ overload during cardiac hypertrophy further enhances cardiac deterioration. Adrenergic catecholaminergic signaling (using isoproterenol) upregulates cardiac contractility, induces greater mitochondrial Ca^2+^ load and oxidative stress. If combined, these events will trigger the mPTP opening (23).

Here, we report that calorie restriction modulates the cellular redox status, leaves mitochondria less sensitive to high ROS levels, and protects mitochondria against damage induced by Ca^2+^ overaccumulation, ultimately leading to anti-hypertrophic effects. Calorie restriction prevented the upregulation of proteins synthesis and the isoproterenol-induced atrial natriuretic peptide (ANP). Our results advance the present knowledge by demonstrating that calorie restriction blocked reverse electron transport-induced high levels of mitochondrial ROS during cardiac hypertrophy. These were accompanied by lower sensitivity to Ca^2+^-induced mitochondrial permeability transition pores. We also demonstrate that calorie-restricted hearts are sensitive to β-adrenergic signaling (stimulated by isoproterenol). Finally, similar to controls, calorie-restricted hearts lose sensitivity after long treatment with isoproterenol. Altogether, our results indicate that short-term calorie restriction protects against cardiac hypertrophy by preserving mitochondrial homeostasis. Additionally, we bring the novel viewpoint of β-adrenergic cardiac stimulation being insensitive to the beneficial effects of calorie restriction in isoproterenol-induced cardiac hypertrophy.

## 2. MATERIALS AND METHODS

### 2.1 Calorie restriction

All animals used were fed with a standard rodent diet. Swiss male mice weighing between 25-30 g were used in experiments involving mitochondrial respiration, H_2_O_2_ production, citrate synthase, and hypertrophy induction. Wistar male rats weighing between 180-240 g were used in hypertrophy induction and isolated cardiac perfusion experiments. For induction of calorie restriction animals were divided randomly into 2 groups: 1. animals fed ad libitum (free access to food – group control) and 2. animals fed with 40% of the ad libitum energy intake (Calorie Restriction group – CR). Animals stayed on this protocol for 24 days. Then, they were randomly divided into 4 other groups and treated with saline or isoproterenol (30 mg/kg per day) for 8 additional days to induce cardiac hypertrophy.

### 2.2 Cardiac Hypertrophy Induction

All procedures were approved by the Institutional Animal Experimentation Ethics Committee of the Universidade Federal do Cariri under protocol number (001/2017). All animals were anesthetized before experiments with ketamine (100 mg/kg of body weight) and xylazine (10 mg/kg of body weight). All experiments were performed according to the Guide for the Care and Use of Laboratory Animals published by the National Institutes of Health. For experiments involving isolated mitochondria, we used 60-day-old Swiss male mice weighing between 25 and 30 g. For experiments involving cardiac Langendorf perfusion, we used male Wistar rats weighing between 180 and 200 g.

Mice fed ad libitum or on a calorie restriction diet were treated daily with isoproterenol intraperitoneally (30 mg/kg/day – ISO group) for 8 days to induce cardiac hypertrophy. For cardiac hypertrophy induction in rats, we injected isoproterenol for 7 days. Animals were sacrificed the next day and injected isoproterenol directly into the heart using a Langendorf perfusion system. The Control group was injected with saline (0.9% - control group). Animals on a calorie restriction diet were first divided into 2 subgroups: one receiving saline (calorie restriction control) and another receiving isoproterenol (30 mg/kg per day – calorie restriction ISO). Cardiac hypertrophy induction was measured by dividing heart weight per tibia length, ventricular atrial natriuretic peptide levels, and by quantifying total cardiac proteins normalized by the wet weight of tissue.

### 2.3 Cardiac langendorff perfusion

Male Wistar rats were sacrificed, the hearts removed and aorta cannulated in no longer than 3 min. Hearts were eliminated from the study if the time between removal and the beginning of perfusion was longer than 3 mins. To retrograde perfuse rat hearts, we used a Langendorff apparatus (AD Instruments) with Krebs-Henseleit buffer containing (in mM) 118 NaCl, 25 NaHCO_3_, 1.2 KH_2_PO_4_, 4.7 KCl, 1.2 MgSO_4_, 1.25 CaCl_2_, 10 glucose, and 10 HEPES, pH 7.4, at 37°C. All perfusions were at a constant flow of 12 mL min^-1^ at 37°C with oxygenated Krebs-Henseleit buffer (gas mixture bubbled with 95% O_2_, 5% CO_2_). The hearts were perfused for 10 minutes in sinus rhythm with oxygenated Krebs buffer before the addition of 50 nM isoproterenol to induce sudden increases in heart rate (chronotropic) and force of contraction (positive inotropic effect). The hemodynamic function was recorded with a fluid-filled latex balloon connected to a pressure transducer (Powerlab/8SP, AD Instruments). This ballon was inserted inside left ventricle. Buffer was supplied to the heart by a peristaltic pump at a constant flow of 12 mL min^-1^.

### 2.4 Cardiac hemodynamic parameters

We continuously monitored the following functional parameters with a computer-based data acquisition system (PowerLab 8S/P with LabChart 4 software, AD Instruments). Left ventricular developed pressure (LVDP) was calculated using the formula LVDP = LVSP-LVEDP where LVSP means left ventricular systolic pressure and LVEDP means left ventricular end-diastolic pressure. We also calculated the time to the peak (TTP) of systolic pressure, pressure-time integral (using 30 seconds fixed time), and heart rate.

### 2.5 Atrial natriuretic peptide detection

Cardiac left ventricles were manually homogenized using a potter in a buffer composed of 300 mM sucrose, 10 mM K^+^ HEPES buffer, pH 7.2, 1 mM K^+^ EGTA, and BSA 1 mg/mL at 4 °C. Nuclei, mitochondria, and cellular residues were pelleted by centrifugation at 9400 g for 5 min. The resulting supernatant was used to assay for ANP levels using a competitive enzyme Immunoassay kit from Sigma-Aldrich (Catalog Number RAB0385). The levels of ANP were normalized for the amount of protein. Results are expressed as relative to controls.

### 2.6 Mitochondrial Isolation

Cardiac tissue was rapidly removed, cut, and washed in a buffer containing 300 mM sucrose, 10 mM K^+^ HEPES buffer, pH 7.2, and 1 mM K^+^ EGTA at 4 °C. Then, protease Type I (Sigma-Aldrich) was used to digest the cardiac tissue for 10 min. The excess protease was removed by washing the cardiac tissue twice with the same buffer containing 1 mg/mL BSA. The tissue was manually homogenized using a glass potter. Nuclei and cellular residues were pelleted by centrifugation at 1200 g for 5 min. To obtain the mitochondrial pellet the supernatant was recentrifuged at 9400 g for 10 min. The pellet was resuspended in a minimal amount of buffer (100 μL) and used at 100 mg/ml for mitochondrial respiration and H_2_O_2_ levels. The supernatant was used for enzymatic assays.

### 2.7 Mitochondrial respiration

We used a Clark-type electrode with constant stirring (“Oxygraph”, “Hansatech”, UK) connected to a data acquisition system to evaluate the mitochondrial oxygen consumption. The reaction media consisted of 150 mM KCl, 10 mM HEPES, 2 mM MgCl_2_, 2 mM KH_2_PO_4_ pH 7.2 (KOH). The respiration trace was initiated with the addition of 0.1 mg of mitochondrial protein. Importantly, the reaction media was allowed to equilibrate in the chamber. State 2 respiration was achieved by the addition of 4 mM succinate. To block the reverse electron transport system the buffer was supplemented with rotenone (1 μM). To test whether our mitochondrial preparation was coupled we conducted a parallel respiratory control ratio experiment by adding 1 mM ADP (state 3), 1 μg/mL oligomycin to induce state 4. To induce the uncoupled state we added 100 nmol CCCP addition. We used mitochondrial preparations with respiratory control ration (state 3/state 4) above 3 (not shown).

### 2.8 Mitochondrial H_2_O_2_

To measure H_2_O_2_ we incubated mitochondria with 4 mM succinate, amplex red (50 μmol/L), and horseradish peroxidase (1 U/mL) for 30 min (in absence of light) at 37°C. The reaction buffer contained: 150 mM KCl, 10 mM HEPES, 2 mM MgCl_2_, 2 mM KH_2_PO_4_ pH 7.2 (KOH). Then, tubes were centrifuged at 9400g for 5 minutes. The supernatant was read at 560 nm in a spectrophotometer zeroed with a blank reaction containing amplex red and horseradish peroxidase without the mitochondrial sample. Mitochondrial H_2_O_2_ production was calculated using H_2_O_2_ standards to create a calibration curve and expressed as μmol/mg protein.

### 2.9 Mitochondrial permeability transition pore (MPTP)

We measure MPTP opening in isolated mitochondria by following the changes in light scattering due to Ca^2+^ uptake. Swelling rates were measured as a decrease over time in optical density at 520 nm. We incubated mouse heart mitochondria (0.2 mg protein/mL) in a buffer containing 100 mM KCl, 10 mM HEPES, 2 mM succinate, 2 mM MgCl_2_, 2 mM KH_2_PO_4_, 1 μg/mL oligomycin, pH 7.2 (KOH). To induce opening of the mitochondrial permeability transition pore, 40 μM Ca^2+^ per mg protein was added.

### 2.10 Citrate synthase activity

Mitochondrial citrate synthase activity was measured by the development of yellow color generated due to the reaction of free Coenzyme A (released from acetyl CoA) with DTNB forming the yellow compound TNB which absorbance is measured spectrophotometrically at 412 nm. The samples were incubated for 5 minutes in a buffer composed of 100 mM Tris HCl pH 8.1, 100 μM DTNB, 300 μM acetyl CoA e 0.1 % Triton X100. Then, 0.1 mM oxaloacetate is added to initiate the reaction. The yellow product was followed by 5 minutes.

### 2.11 Superoxide Dismutase (SOD) assay

A buffer containing: 0.1 mM EDTA, 13 mM L-methionine, and 75 mM nitro blue tetrazolium (NBT) in potassium phosphate buffer (pH 7.8) was used to measure SOD activity. Just after the addition of cardiac homogenates 2 μM riboflavin was added. The samples were exposed uniformly to an unfiltered white light for 10 minutes. The developed blue color due to NBT reduction was measured at 560 nm. SOD activity was expressed as U/mg of protein. One unit is the amount of enzyme required to inhibit the reduction of NBT by 50%. The enzymatic activity in each sample was correlated with the gross indicator of cardiac hypertrophy heart weight per tibia length.

### 2.12 Glutathione peroxidase activity assay

Glutathione peroxidase activity was determined in the presence of 1 mM cumene hydroperoxide (Sigma-Aldrich) as a substrate. The activity was assayed by the decrease in NADPH. A buffer containing: 50 mM potassium phosphate, 0.5 mM EDTA, 0.2 mM NADPH (Sigma Aldrich), 1 mM GSH (Sigma-Aldrich), and 0.2 U/mL glutathione reductase purified from S. cerevisiae (Sigma Aldrich), pH 7.0 was used. The cellular extract was incubated in this buffer for 5 min. Then, cumene hydroperoxide was added to initiate the reaction. We followed the disappearance of the co-substrate NADPH at 340 nm for 5 min. Glutathione peroxidase activity is expressed as Units/mg protein. The enzymatic activity in each sample was correlated with the gross indicator of cardiac hypertrophy heart weight per tibia length.

### 2.13 Statistical Analysis

Data are presented as mean±SEM. Comparisons between groups were conducted one-way ANOVA followed by a Tukey’s test. P < 0.05 was considered to be statistically significant. All vector graphics were created using Graphpad prism software. Associations between heart weight/tibia length, and antioxidant enzymes activities (superoxide dismutase and glutathione peroxidase) were tested for significance using Pearson’s correlation coefficients.

## 3 RESULTS

### 3.1 Calorie restriction protects mice from developing cardiac hypertrophy

To demonstrate the effects of calorie restriction on cardiac hypertrophy, we subjected mice to isoproterenol (30 mg/kg/day) for eight days. Control mice treated with isoproterenol showed a significant increase in the gross indicator of cardiac hypertrophy heart weight per tibia length (HW/TL) and increased protein levels per gram of tissue. Short-term calorie restriction blocked these hypertrophic gross indicators markers (Fig. 1A, B). The hypertrophic stimulus causes the translocation of the nuclear factor of activated T-cells (NFAT) transcription factor to the nucleus resulting in the activation of the fetal gene program - indicated by higher cardiac ANP levels (24, 25). To test whether calorie restriction can block the increase in ANP levels, we detected ANP directly into the heart homogenate by ELISA. Calorie restriction resulted in a marked decrease in hypertrophic ANP levels (Fig. 1C). These results confirm previous findings from our group (1) and others (10, 26) that calorie restriction is a powerful antihypertrophic procedure.

**FIGURE 1.**
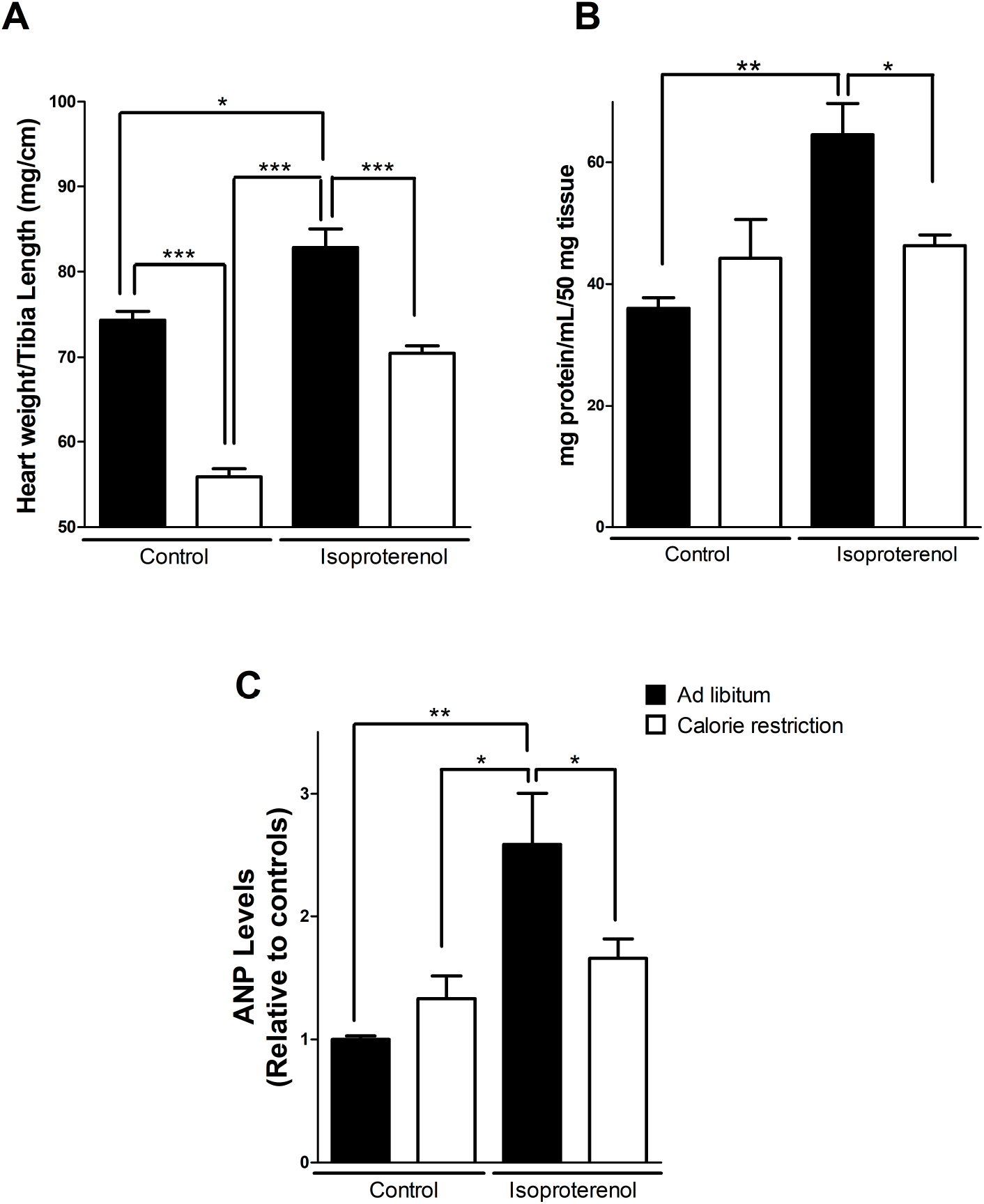
Short-term calorie restriction blocks isoproterenol-induced cardiac hypertrophy. A. Heart weight/tibia length analysis of cardiac tissue from controls or calorie-restricted mice treated or not with isoproterenol (30 mg/kg/day). B. Quantification of cardiac protein. Statistical significance was determined by one-way ANOVA followed by a Tukey post hoc test. P **, P < 0.01. P ***, P < 0.001.

### 3.2 Calorie restriction blocks mitochondrial reverse electron transport-derived H_2_O_2_ accumulation during cardiac hypertrophy

A large body of evidence has implicated ROS generation in cardiac hypertrophy/heart failure. Antioxidant enzyme repression is also a hallmark of cardiac hypertrophy (for review see, (21)). The oxidative stress generated during cardiac hypertrophy could result from a dysfunctional respiratory chain. Indeed, we found that mitochondrial complex I activity is suppressed during isoproterenol-induced cardiac hypertrophy (Results not shown). Here, we investigated whether cardiac hypertrophy enhances mitochondrial ROS production by reverse electron transport. To accomplish this, we energized mitochondria with succinate (complex II substrate) in the presence or absence of rotenone. Rotenone in these conditions blocks the mitochondrial reverse electron transport. Then we probed for H_2_O_2_ production using Amplex Red for 30 min at 37 °C (Fig. 2A). Fig 2A also brings a representative oxygen consumption trace. Fig. 2B shows that mitochondria isolated from hypertrophic samples, energized as described above, released significantly higher amounts of H_2_O_2_. As expected, rotenone (a complex I inhibitor) blocked it. Strikingly, calorie restriction avoids H_2_O_2_ production by reverse electron transport. Surprisingly, the H_2_O_2_ levels in calorie restriction are insensitive to rotenone suggesting that reverse electron transport is not the source of H_2_O_2_ seen in these samples or that calorie restriction induces a higher antioxidant system which could block mitochondrial H_2_O_2_ release. Interestingly, respiratory rates (Fig. 2C) in mitochondria energized with succinate or succinate plus rotenone are equal. To further understand the mitochondrial effects of calorie restriction during isoproterenol-induced cardiac hypertrophy, we determined the activity of citrate synthase, a mitochondrial matrix enzyme often used as a marker of mitochondrial content. Surprisingly, hypertrophic samples had preserved citrate synthase activity. Calorie restriction significantly enhanced it during hypertrophic insult (Fig. 3). These results suggest that calorie restriction avoids cardiac hypertrophy impacting mitochondria and oxidative stress.

**FIGURE 2.**
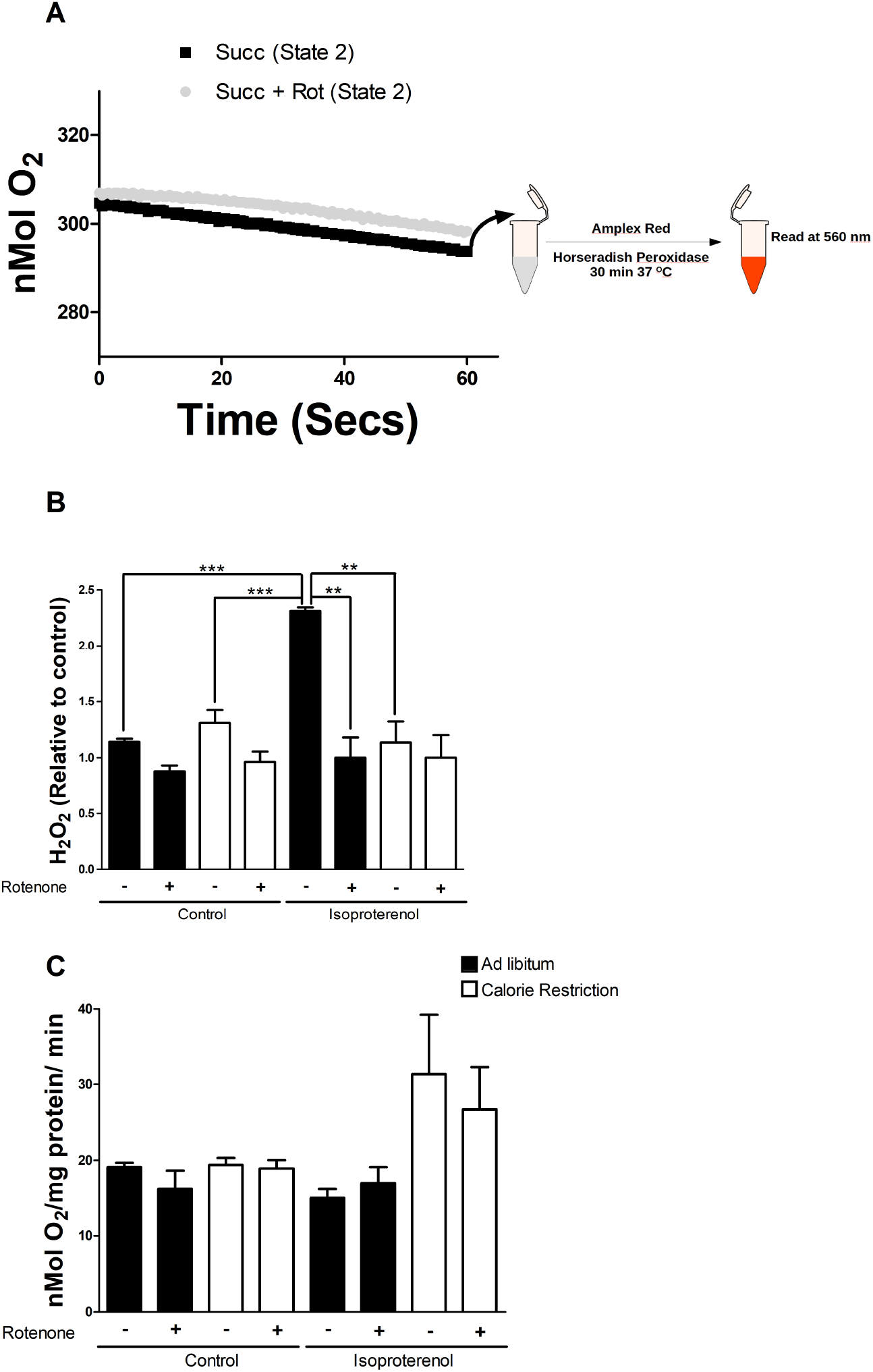
Calorie restriction avoids H_2_O_2_ production by mitochondrial reverse electron transport during cardiac hypertrophy. Mitochondria (0.1 μg protein/mL) were incubated in 150 mM KCl, 10 mM HEPES, 4 mM succinate, 2 mM MgCl_2_, 2 mM KH_2_PO_4_, pH 7.2 (KOH). **A.** Representative trace of mitochondrial respiration in each condition. Scheme of samples collection and H_2_O_2_ detection. **B.** Represent average±standard errors of mitochondrial H_2_O_2_ production. Rotenone (1 mM) was added to block mitochondrial reverse electron transport. **C.** Quantification of mitochondrial oxygen consumption in each condition. Graphs represent 3 or 4 different experiments in each condition. Statistical significance was determined by one-way ANOVA followed by a Tukey post hoc test. P **, P < 0.01. P ***, P < 0.001.

**FIGURE 3.**
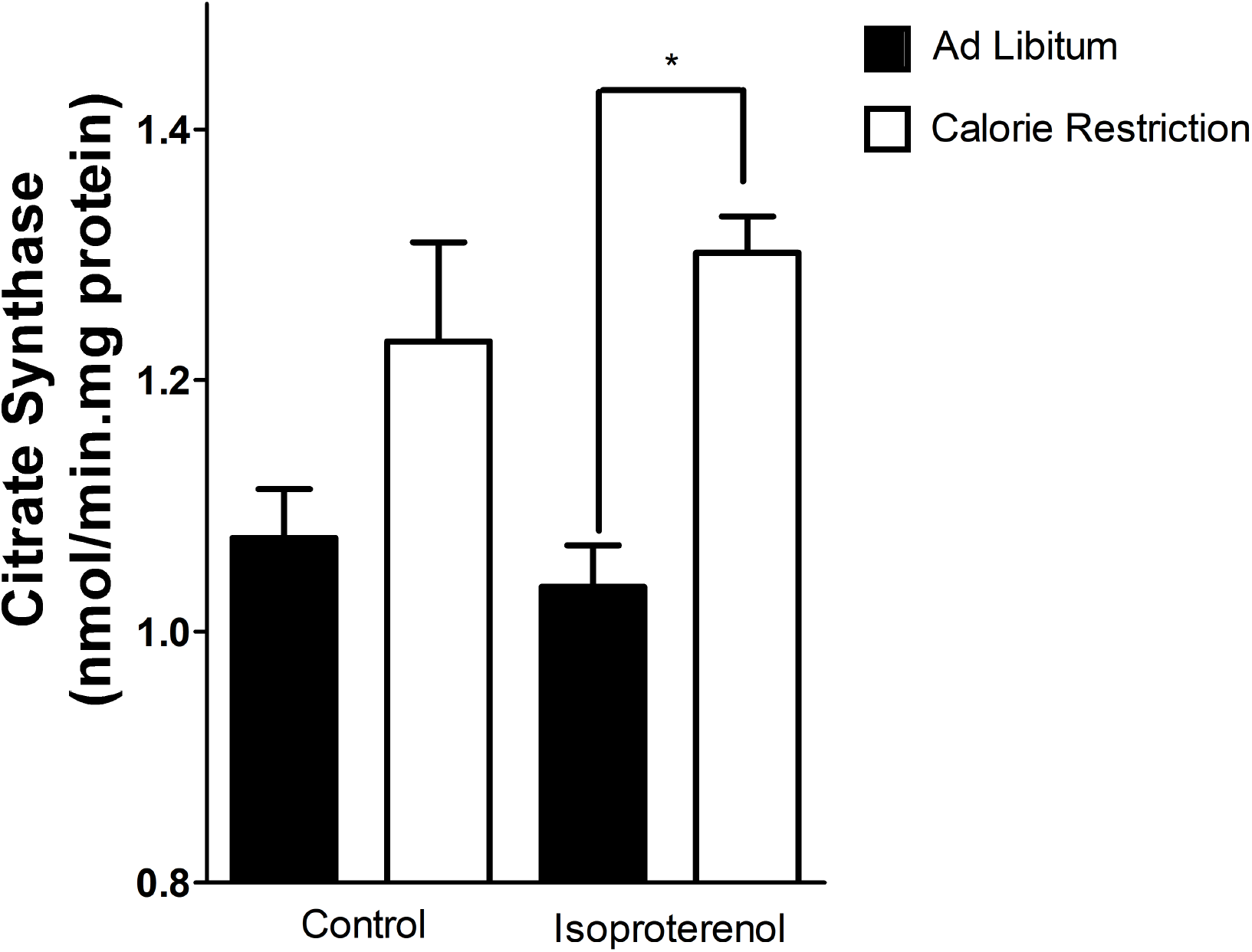
Calorie restriction enhances mitochondrial content during isoproterenol-induced cardiac hypertrophy. Mitochondrial citrate synthase activity was measured spectrophotometrically at 412 nm by the formation of TNB as described in materials and methods. The graph represents 4 different experiments in each condition. Statistical significance was determined by one-way ANOVA followed by a Tukey post hoc test. *, P < 0.05.

We then asked if calorie restriction affected cardiac antioxidant enzymes. Indeed, we have shown that cardiac hypertrophy is powerful enough to avoid antioxidant enzymes activity repression during cardiac hypertrophy (1). Here, we took another approach by running a correlation analysis between superoxide dismutase and glutathione peroxidase activity and heart weight per tibia length (a gross indicator of cardiac hypertrophy). Strikingly, we found a highly significant negative correlation between heart weight/tibia length and both enzyme activities (Fig. 4A, B). These results suggest that calorie restriction blocks oxidative stress during cardiac hypertrophy, limiting cardiac pathological growth.

**FIGURE 4.**
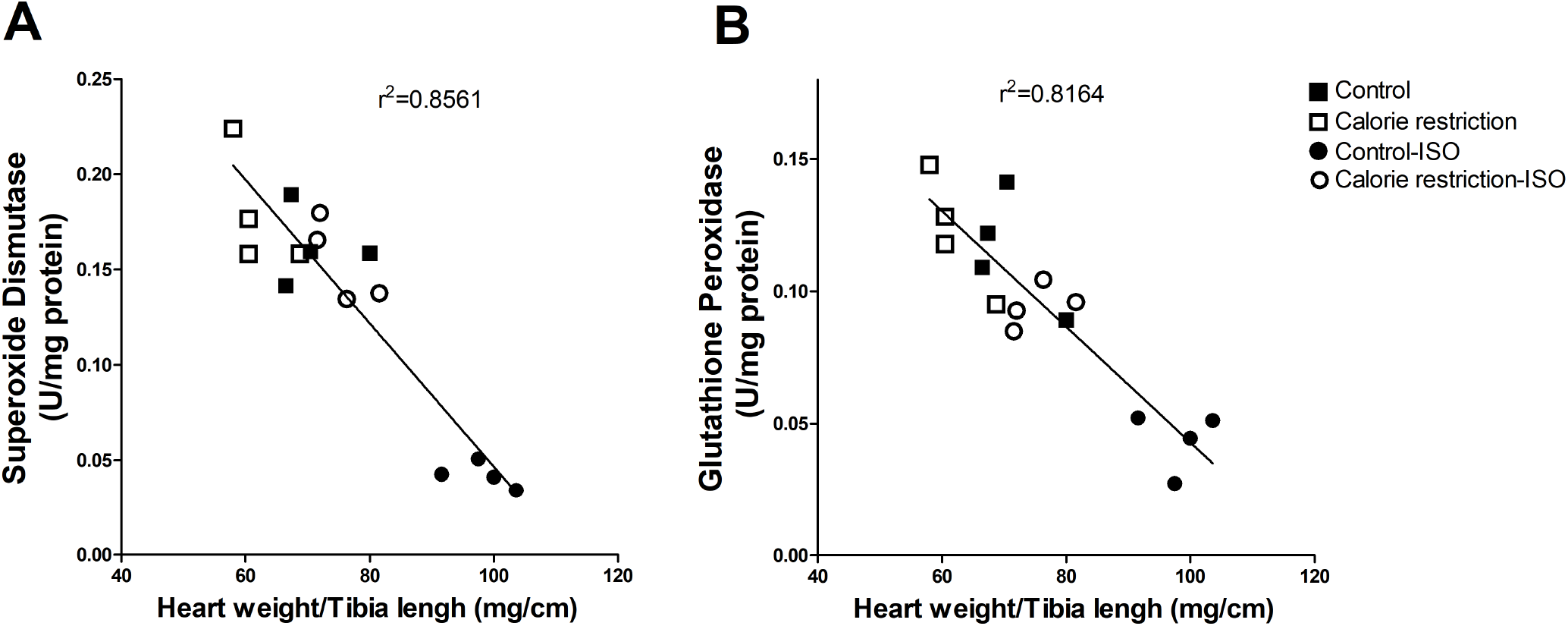
Cardiac hypertrophy indicated by HW/TL negatively correlates with antioxidant enzyme activity. **A.** Correlation of cardiac hypertrophy (HW/TL) with superoxide dismutase activity (A) and glutathione peroxidase activity (B). ISO = isoproterenol 30 mg/Kg/day.

### 3.3 Isolated mitochondria from calorie-restricted mice are protected against calcium-induced mitochondrial permeability transition pore

Mitochondria from hypertrophied hearts are vulnerable to stress-induced (by Ca^2+^ overload and oxidative stress) opening of the mitochondrial permeability transition pore (mPTP) (23, 27). To investigate whether calorie restriction protects against mPTP, we treated isolated cardiac mitochondria (energized as Fig. 2) with Ca^2+^ (2 μM/0.05 mg mitochondrial protein) in the presence of succinate with no rotenone (a condition which generates high reactive oxygen species). Mitochondria from calorie-restricted mice had no difference in susceptibility to mPTP when compared to control samples. On the other hand, isolated mitochondria from hypertrophic hearts had significantly higher susceptibility to Ca^2+^-induced mPTP opening. Indeed, these samples had a greater decrease in light scattering in 540 nm. Calorie restriction completely reversed this effect (Fig. 5). Thus, calorie restriction represents a non-pharmacological tool that protect mitochondria against Ca^2+^-induced mPTP.

**FIGURE 5.**
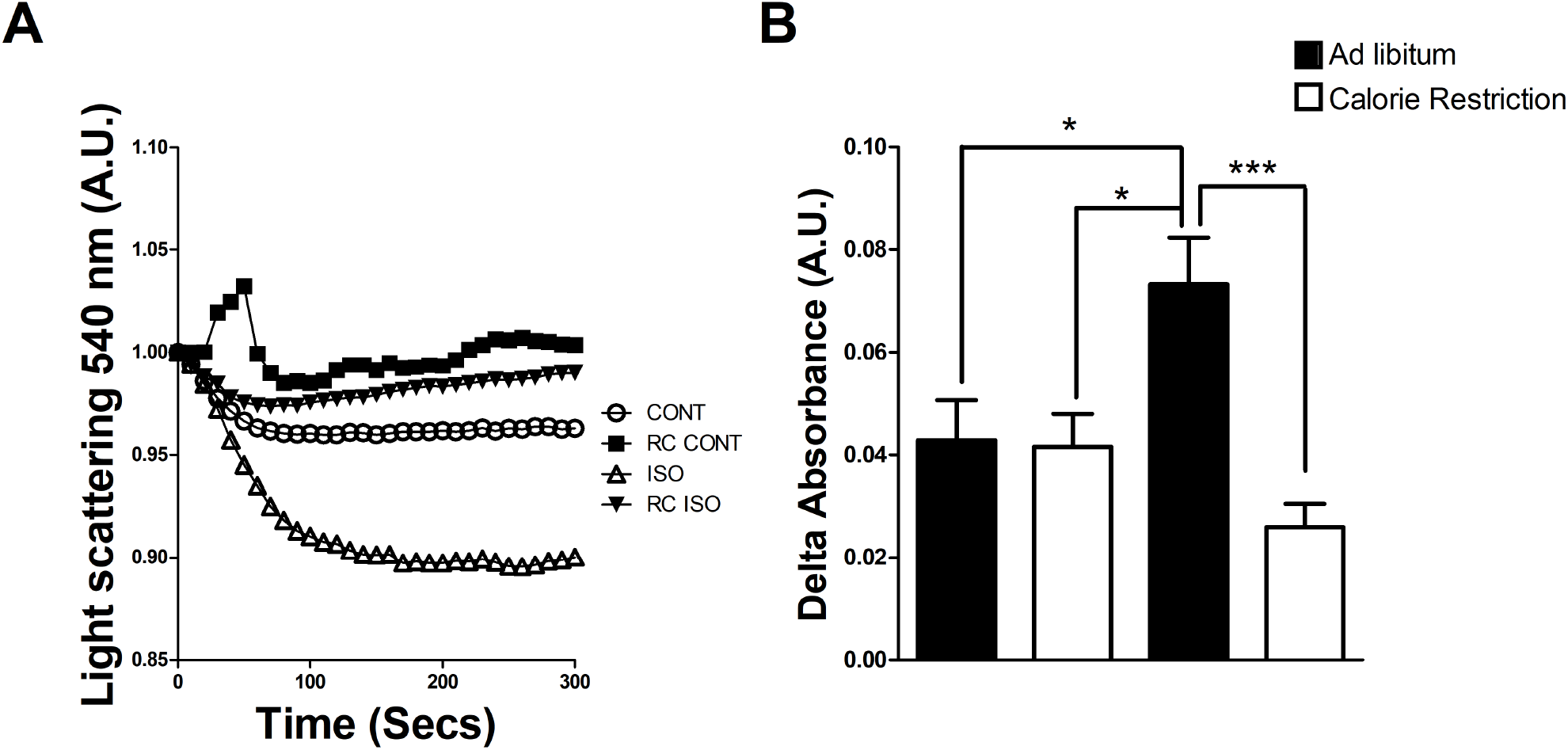
Calorie restriction protects mitochondria against Ca^2+^-induced swelling. Cardiac mitochondria were isolated from controls and calorie-restricted mice treated or not with isoproterenol. Mitochondria (0.1 μg protein/mL) were incubated in 100 mM KCl, 10 mM HEPES, 4 mM succinate, 2 mM MgCl_2_, 2 mM KH_2_PO_4_, 1 μg/mL oligomycin, pH 7.2. Mitochondrial permeability transition pore opening was induced by the addition of 2 μM Ca^2+^. A. representative traces of 6 independent experiments. B. Delta of absorbance (540 nm) between 5 and 0 minutes time point for each curve. *, P < 0.05, ***P < 0.001.

### 3.4 Catecholamine signaling is preserved in calorie-restricted hearts

One could argue that calorie restriction renders mice insensitive to β-adrenergic signaling. In this hypothesis, hearts from calorie-restricted mice would be less hypertrophic due to a lower induction. To test this, we treated isolated rat hearts with 50 nM isoproterenol in a Langendorf apparatus (scheme of treatment, Fig. 6A). As expected, the heart rate from calorie-restricted mice in sinus rhythm was similar to controls (Fig. 6B). It is important to note that isoproterenol (applied directly to the heart) stimulated the pacemaker cells, significantly enhancing the heart rate in both groups (Fig. 6B). Isoproterenol non-specifically activates β1 receptors, which results in PKA-dependent positive inotropic and chronotropic responses (28). To investigate isoproterenol-induced inotropic effects, we quantified the cardiac left ventricular developed pressure (LVDP, Fig. 6C) and pressure-time integral (Fig. 6D) in the Langendorf apparatus. At baseline, both measurements were similar between the groups. It is important to note that hearts treated with isoproterenol (50 nM) had increased LVDP and pressure time integral in both groups. No significant differences were seen in time to the peak pressure between controls and calorie-restricted mice (Fig. 6D). These results suggest that calorie restriction has preserved β1 receptors signaling (chronotropic and inotropic effects). These results exclude any possible downregulation of catecholaminergic signaling in control calorie-restricted hearts. Therefore, we sustain that mitochondrial and redox changes are responsible for the antihypertrophic effects of calorie restriction.

**FIGURE 6.**
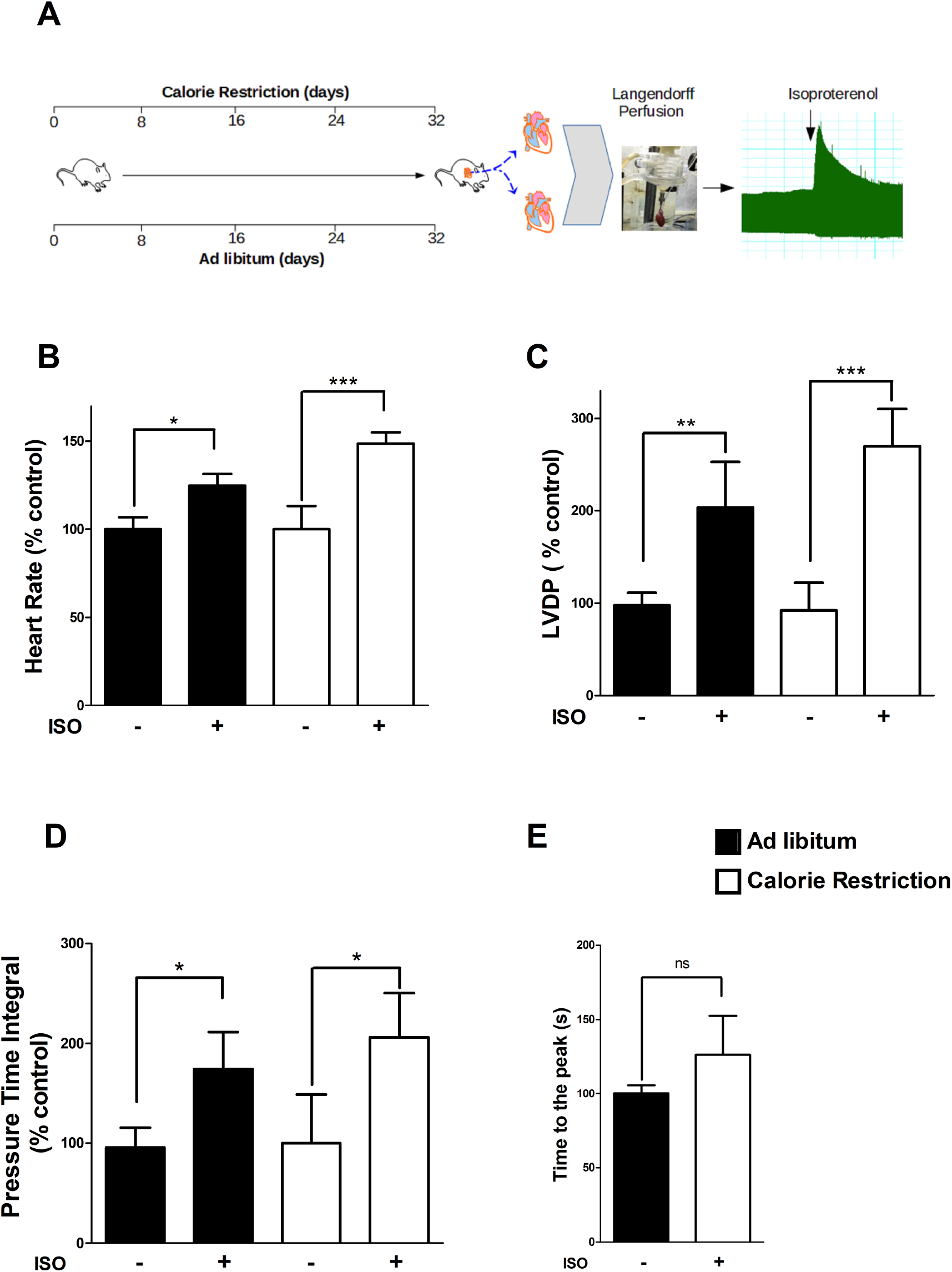
Cathecolamine signaling is preserved in calorie-restricted hearts. Isolated perfused rat hearts were stimulated with 50 nM isoproterenol. Heart rate (A), left ventricular developed pressure (LVDP, B), pressure-time integral (C), and time to the peak (D) were measured before and after the beta-adrenergic stimulus of 50 nM isoproterenol. The graphs represent 4 different experiments in each condition. Statistical significance was determined by one-way ANOVA followed by a Tukey post hoc test. *, P < 0.05, **, P < 0.01, P ***, P < 0.001.

### 3.5 Isoproterenol-induced cardiac hypertrophy blocks cardiac catecholaminergic signaling

Isoproterenol (a non-selective β-agonist) binds cardiac beta receptors resulting in increased positive inotropic and chronotropic responses. Knowing that β1 catecholaminergic signaling is preserved during calorie restriction we tested the effects of isoproterenol (50 nM) on rat hearts treated with isoproterenol for seven days (30 mg/Kg/day – scheme in Fig. 7A). Isoproterenol treatment (7 days) in control rats resulted in massive cardiac hypertrophy as estimated by heart weight per tibia length (HW/TL) ratio (Fig. Supplementary 1). Interestingly, cardiac rate (Fig. 7B), left ventricular developed pressure (Fig. 7C), and pressure-time integral (Fig. 7D) were not affected by ex vivo isoproterenol infusion. Several groups have reported a reduced inotropic response to beta-adrenergic stimulation during cardiac hypertrophy (29–32). Surprisingly, we also found reduced isoproterenol-induced inotropic and chronotropic effects in calorie-restricted hearts (Fig. 7B, C, D, and E). Finally, we also found no difference in time to the peak between both groups. We believe these effects might be due to a reduced β-adrenergic signaling. A possibility exists that would be a result of poor mitochondrial performance. We do not exclude a poor mitochondrial performance in isoproterenol-treated calorie-restricted hearts, but we have evidence for preserved or slightly enhanced mitochondrial oxygen consumption in Fig. 2.

**FIGURE 7.**
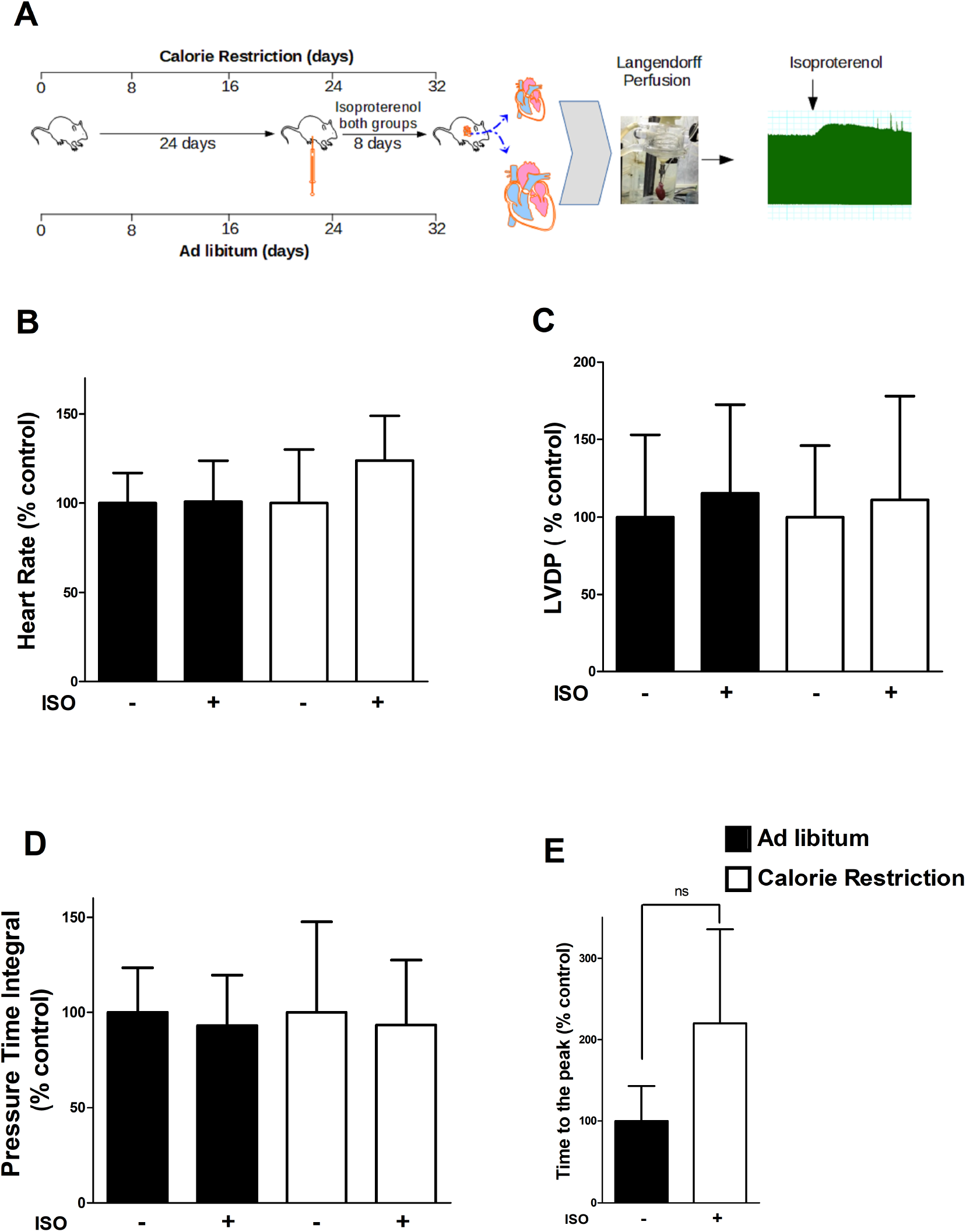
Cathecolamine signaling is suppressed in isoproterenol-treated hearts. Isolated perfused rat hearts were stimulated with 50 nM isoproterenol. Heart rate (A), left ventricular developed pressure (LVDP, B), pressure-time integral (C), and time to the peak (D) were measured before and after the β-adrenergic stimulus of 50 nM isoproterenol. The graphs represent 4 different experiments in each condition. Statistical significance was determined by one-way ANOVA followed by a Tukey post hoc test. *, P < 0.05, **, P < 0.01, P ***, P < 0.001.

## 4 DISCUSSION

By combining isolated heart and mitochondrial experiments we show that short-term calorie restriction applied to mice is a powerful anti-hypertrophic tool. The data indicate a robust and powerful reversal of hypertrophic oxidative stress by calorie restriction during β-adrenergic-stimulated cardiac remodeling. This is particularly seen when we forced more reactive oxygen species production by energizing mitochondria with succinate in the absence of rotenone (a complex I blocker). Mitochondria energized with succinate produces higher amounts of H_2_O_2_ (by reverse electron transport). Mitochondria isolated from hypertrophic hearts had the highest succinate-induced H_2_O_2_ levels. Remarkably, calorie restriction prevents reverse electron transport-induced ROS generation and enhances antioxidant enzymes. We also demonstrate that mitochondria isolated from calorie restriction hearts are protected against the Ca^2+^-induced opening of the mitochondrial permeability transition pore. Our model involves subjecting mice to isoproterenol-induced catecholaminergic signaling to induce cardiac pathological growth. One could argue that calorie-restricted mice were resistant to isoproterenol at baseline. To answer this, we show that calorie-restricted hearts have preserved isoproterenol-induced chronotropic and inotropic effects (Fig. 6). It is important to note that the cardiac tissue of mice treated with isoproterenol for seven days was insensitive to this drug in the Langendorf system (Fig. 8). Finally, we summarize the findings of this paper in Fig. 8 where we conducted a multivariate analysis using the metaboanalyst website. We analyzed the data from SOD, GPX, H_2_O_2_, cardiac ANP, and protein levels using principal component analysis (PCA) analysis which showed a distinct separation of the cardiac hypertrophic group and all the other groups (Fig. 8B). Using the patternhunter test, we found that ANP and H_2_O_2_ positively correlate with cardiac protein levels. Of further notice, we found a negative correlation between both SOD and GPX activities and cardiac protein levels (Fig. 8C). To the best of our knowledge, this is the first report describing the beneficial effects of calorie restriction on mitochondrial reactive oxygen species production by reverse electron transport, on the opening of the mitochondrial permeability transition pore, and the sensitivity of calorie-restricted hearts to catecholamines.

**FIGURE 8.**
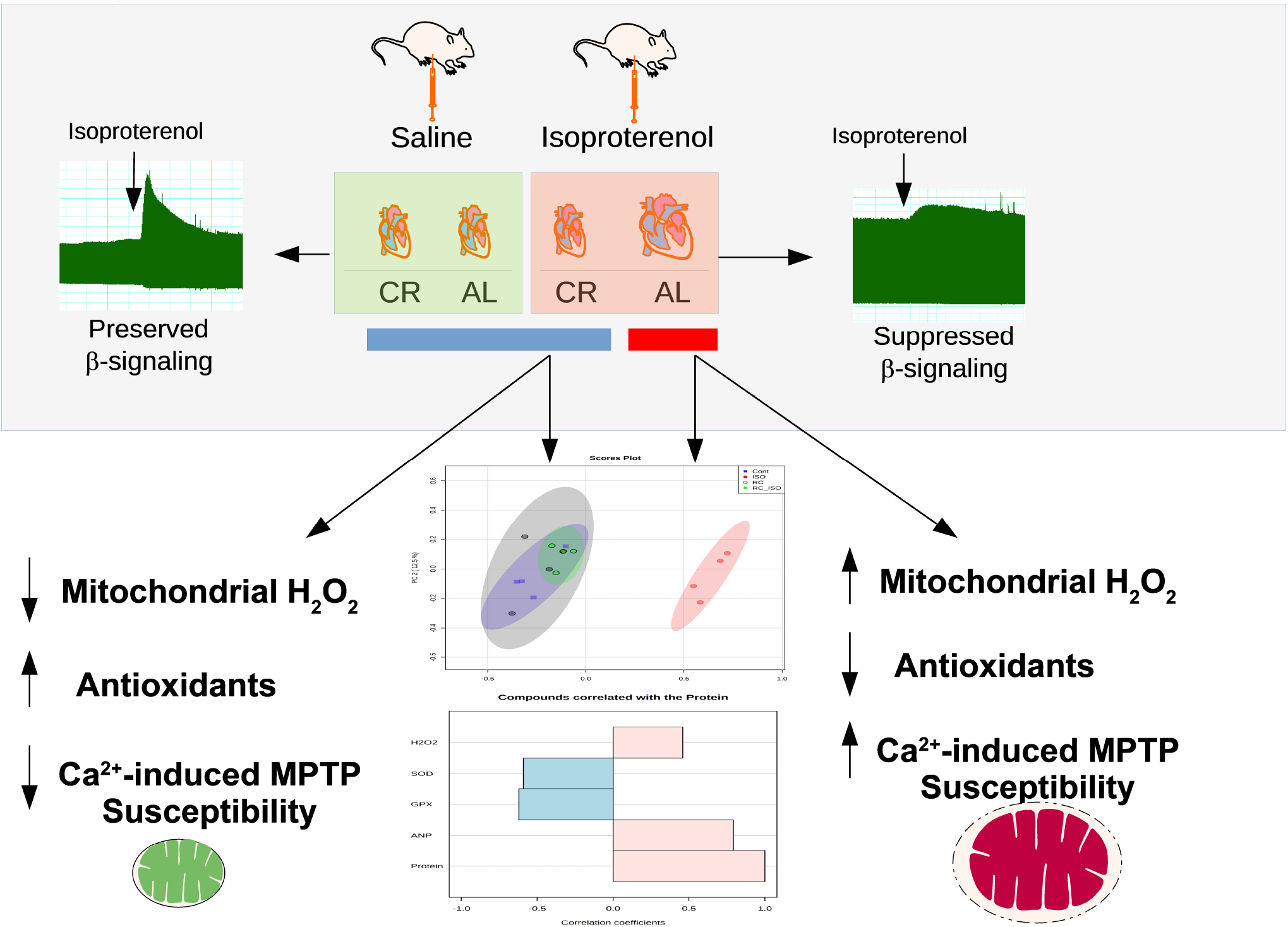
Summary of main paper findings. Scheme showing the main findings of the paper. The top panel shows the suppression in β-adrenergic stimulation by isoproterenol in hypertrophic samples. The bottom panel shows that calorie restriction blocks reactive oxygen species (H_2_O_2_) elevation, increases antioxidants, and avoids swelling secondary to Ca^2+^-induced mitochondrial permeability transition pore opening. The bottom panel shows principal component analysis (top graphic) and patternhunter test (bottom graphic) of SOD, GPX, H_2_O_2_, cardiac ANP, and cardiac protein levels.

Dysregulated catecholaminergic and adrenergic signaling are risk factors for hypertrophy and heart failure development (20). Indeed, some studies reported a reduced response to beta-adrenergic stimulation during cardiac hypertrophy (29–32). Our data confirm a reduction in cardiac beta-adrenergic stimulation capacity during cardiac hypertrophy. Surprisingly, calorie-restricted hearts (treated for seven days with isoproterenol) had decreased beta-adrenergic response despite blocking several markers and consequences of cardiac hypertrophy. This result suggests that β-adrenergic signaling suppression may be disconnected from the recovered phenotype. It is important to note that our calorie restriction procedure is short-term, which could reverse several of the signals and alterations present in cardiac hypertrophy but not all of them. Indeed, a long-term calorie restriction procedure improved cardiac inotropic reserve and sympathetic innervation in an animal model of ischemia-induced heart failure (33). Another possibility is that β-adrenergic signaling suppression is a protective feedback mechanism in both controls and calorie restrict animals (34). It is important to note that β-blockade attenuates dysfunction in infarcted hearts (35). Long-term exposure to catecholamines induces desensitization of receptors (36). Evidence points to a desensitized cardiac β-adrenergic signaling (with preserved catecholamine levels) during cardiac hypertrophy (31). We believe that the downregulation of β-adrenergic signaling would help counteract the pathogenic mechanisms of hyperactivation of the sympathetic nervous system (especially this is true during heart failure) (37).

Mitochondria is a major source of ROS inside cardiomyocytes. These cells are packed with mitochondria which supply the majority of the ATP necessary for cardiac contractility. On the flipside, mitochondria may generate high levels of ROS, especially in pathological conditions, leading to cellular damage (for review, please see (38)). The imbalance between ROS generation and its clearance by antioxidants results in oxidative stress, which is associated with cardiac hypertrophy conditions (5, 39–42). Here, we simulated impaired mitochondrial ROS formation by energizing isolated mitochondria with succinate with no rotenone (reverse electron transport). This condition will generate high mitochondrial ROS in a manner blocked by rotenone (a complex I inhibitor). This condition is especially important after sustained cardiac ischemia that drives high levels of succinate accumulation. Accumulated succinate will induce mitochondrial reverse electron transport and ROS production during reperfusion (43). Using this approach we found that mitochondria from hypertrophic hearts generated higher levels of ROS than controls. Notably, these changes were significantly blocked by calorie restriction. Interestingly, we found no differences between mitochondrial H_2_O_2_ production of controls or calorie-restricted mice with no isoproterenol treatment. Others have found that calorie restriction significantly decreased mitochondrial ROS levels (44). Additionally, this was accompanied by a lower state 3 (ADP added) oxygen consumption. Mixed results have also been reported previously with some studies reporting unaltered or decreased reactive oxygen species release in response to CR (16). Importantly, we found that succinate-driven mitochondrial oxygen consumption was similar among all groups in state 2. One important observation is that all other papers induced calorie restriction for several months. Our protocol is just for a month, and the basal level of ROS might change in a much more advanced time point. Most importantly, and as said before, our protocol of calorie restriction is sufficiently robust to completely block the reverse electron transport-induced mitochondrial ROS release during cardiac hypertrophy. We have seen similar results when probing for tissue-released H_2_O_2_ (1). Indeed, hypertrophied hearts have higher oxidative stress (5, 39–42). Tissue oxidative stress also depends on the cardiac levels of antioxidants. This scavenging system would influence the maximal ROS release. Isolated mitochondrial studies are highly informative by eliminating or approximating the maximal ROS seen *in vivo* with no influence of cytosolic antioxidant systems. Here, we report higher levels of H_2_O_2_ release by isolated mitochondria that are sensitive to calorie restriction. Besides this observation, we provide data showing that cardiac hypertrophy is accompanied by SOD and GPX repression with a strong negative correlation between SOD or GPX and the gross hypertrophic indicator heart weight per tibia length (Fig. 4). Repression of antioxidant enzymes and higher oxidant production occur in cardiac hypertrophy (40, 42, 45). Blocking higher oxidant production by enhanced antioxidant status avoids cardiac hypertrophy (5, 39, 46). In this sense, calorie restriction avoids cellular antioxidant repression leading to anti-hypertrophic effects. These effects occur together with lower mitochondrial ROS production.

Besides changing mitochondrial/cellular redox susceptibility, we found that calorie restriction blocks Ca^2+^-induced MPTP in isolated mitochondria. MPTP is stimulated by mitochondrial overaccumulation of Ca^2+^, protein thiol oxidation, and high levels of oxidants (22). Isoproterenol treatment upregulates mitochondrial Ca^2+^ accumulation, which also leads to oxidative imbalance besides inducing higher inotropic and chronotropic effects (23). A possibility exists that calorie restriction could avoid MPTP opening by decreasing mitochondrial Ca^2+^ uptake. Nevertheless, recent studies have shown that calorie restriction does not alter the mitochondrial Ca^2+^ transport (44). In this study, we reported the susceptibility of mitochondria to MPTP opening on cardiac samples. Calorie restriction blocked the Ca^2+^-induced swelling (MPTP) on isoproterenol-treated samples. We believe that some factors contribute to this. Specifically, the lower ROS generation in calorie restriction samples, seen in Fig. 2, and preserved oxidant scavenging protect mitochondria by decreasing the probability for Ca^2+^-induced MPTP opening. These data strongly suggest that calorie restriction is a protective dietetic procedure leading to the idea that MPTP and mitochondrial damage may be modulated by calorie restriction.

## 5. CONCLUSION

Taken together, these results suggest, through a series of mitochondrial, oxidative stress, and cardiac hemodynamic studies, that calorie restriction possesses beneficial effects against hypertrophic cardiomyopathy. However, it may lack effects on some of the hypertrophic consequences, such as β-adrenergic signaling repression.

## ACKNOWLEDGEMENTS

The authors acknowledge the technical assistance of Anna Lidia Nunes Varela, Iuliana Marjory Martins Ribeiro, and Antônio F. R. Santos. The authors would like to thank Dr. Marcus Oliveira for his helpful discussions and critical evaluation of the manuscript.

## AUTHORSHIP CONTRIBUTION

Study concept and design: Heberty T Facundo; Aline Maria Brito Lucas; Data acquisition: Plinio Bezerra Palacio, Pedro Lourenzo Oliveira Cunha, Aline Maria Brito Lucas; Data analysis: Aline Maria Brito Lucas and Heberty T Facundo. Heberty T Facundo wrote the paper, which was critically reviewed by all authors. All authors read and approved the final manuscript.

