## Supplementary figures and images for "Effects of calorie restriction on reactive oxygen species production by mitochondrial reverse electron transport, mitochondrial permeability transition pore, and beta-adrenergic stimulation during cardiac hypertrophy"

### Supplemental Figure 1

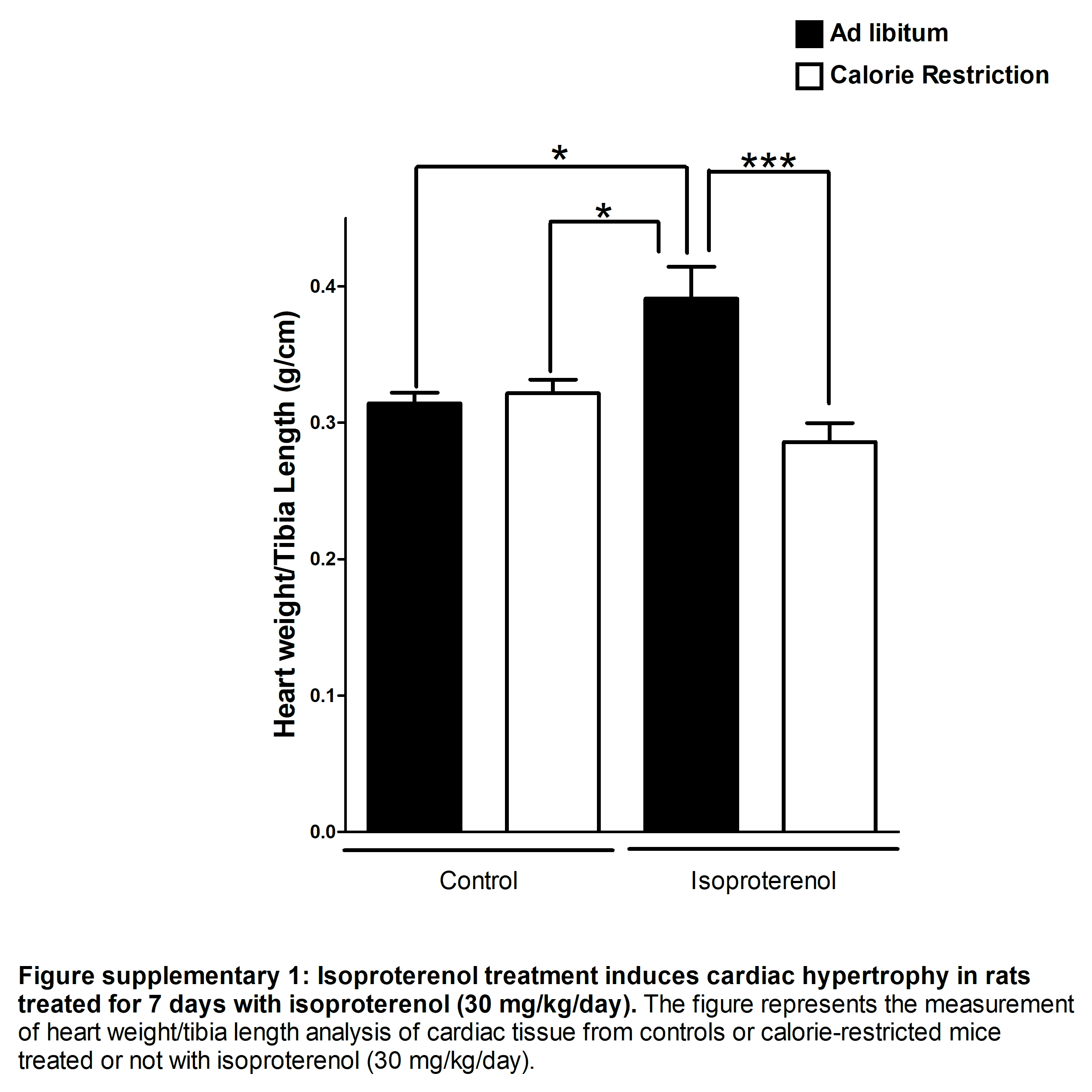
